# Mindin mediated αM-Integrin endocytosis activates STAT3 to maintain keratinocyte stemness

**DOI:** 10.1101/2025.08.24.669246

**Authors:** Binita Dam, Johan Ajnabi, Tirthankar Saha, Ashish Shrivastava, Krithika Badarinathan, Akshay Hegde, Abhik Dutta, Siddham Jasoria, Sunny Kataria, Ashutosh Singh, Colin Jamora

## Abstract

Signalling networks rely not only on specific protein interactions but also on the subcellular locales at which these occur. Such spatial control is fundamental to cellular behaviour in both physiological and pathological scenarios. In studying keratinocyte stemness, we previously identified a Mindin–αMβ2 (CD11b/CD18) interaction that triggers STAT3 activation. Here, we delineate the mechanism underlying this response and show that the F-Spondin domain of Mindin constitutes the minimal integrin-binding module required to initiate downstream signalling. F-Spondin binding to the integrin at the plasma membrane does not elicit the full activation state of the integrin. Instead, it promotes Src-kinase dependent endocytosis of the integrin receptor to the early endosomes. Analysis of integrin conformational dynamics reveals that this compartmental shift is essential as the acidic environment of early endosomes drives the transition into a signalling-competent state. This mechanism extends to pathological contexts, as we demonstrate a requirement for endocytosis in activating STAT3 signalling and preserving stem-like properties in a cancer stem cell model. Together, these findings highlight a previously unrecognized layer of spatial control in integrin signalling, confirming endosomal trafficking as a critical determinant of stem cell behaviour and offering new conceptual and therapeutic opportunities across regenerative biology and cancer.

## INTRODUCTION

Cellular behaviour in homeostasis and disease is governed by tightly regulated signalling networks that integrate extracellular cues into intracellular responses controlling cell fate, survival, and function. These responses are shaped both by how signalling proteins physically interact and by the spatial and temporal context of their activation. Spatial regulation is mediated by endomembrane compartments including the ER, Golgi, endosomes, and lysosomes which function as active signalling platforms beyond their canonical trafficking roles. (*1*, *2*). This raises the question of why the cell uses endomembrane spaces as signalling hubs.

A wide range of ligand-receptor systems illustrate that endomembrane modify their signalling outcomes. G-protein coupled receptors (GPCRs) activated at the cell surface can continue their downstream signalling from the Golgi to generate compartment-specific G-protein and cAMP responses (*1*). Transforming growth factor-β (TGFβ) receptors most effectively activate Smad2/3 in early endosomes. Receptor tyrosine kinases (RTKs) such as epidermal growth factor receptor (EGFR) sustains MAPK and Akt pathways from early endosomes, while their trafficking to late endosomes and lysosomes attenuates signalling through degradation (*2–6*). Integrins also undergo continuous cycles of endocytosis and recycling (*7*), and emerging evidence shows that internalized integrins can continue to signal from endosomal compartments (*8*). These suggest that the endomembrane compartments provide a scaffold where signals are initiated, amplified, or terminated.

This leads to the important question of how do these compartments regulate signalling events. A prominent example is the endosomal control of the transcription factor signal transducer and activator of transcription factor 3 (STAT3) activation by cytokines of the interleukin-6 (IL-6) family. IL-6-bound receptor is activated at the plasma membrane and subsequently trafficked into the endosomal system, where internalised receptor complexes phosphorylate STAT3 in early and late endosomes. These intracellular compartments carrying activated receptors helps in recurrent activation of STAT3 and shifts the balance between nuclear import and export of STAT3 toward sustained import, resulting in nuclear enrichment of STAT3. This endosomal mode of STAT3 activation produces prolonged transcriptional responses essential for key cellular functions (*9*, *10*). One mechanism contributing to this endosomal regulation has been described in embryonic and hematopoietic stem cells, where the endosomal adaptor OCIA interacts with STAT3 and directs it to endosomes, thereby supporting efficient activation by internalised IL-6 receptor complexes (*10*). Notably, aberrant spatial regulation of signalling networks through mis-localised receptors, trafficking defects, or altered endomembrane dynamics has been increasingly linked to disease progression (*11*, *12*). Therefore, understanding how spatial organisation shapes signalling in physiology is critical for explaining how its disruption contributes to pathological conditions. Recently, we identified Mindin (Spondin-2), a soluble matricellular protein, as a novel, non-canonical regulator of STAT3 activation to maintain the stem/progenitor state of epidermal keratinocytes stem cells. Mindin activates STAT3 through its interaction with ɑM-integrin (CD11b) at the surface of keratinocytes. Disruption of this interaction abrogates the activation of the STAT3 pathway and keratinocytes lose their stemness characteristics (*13*). However, the precise mechanism by which the Mindin–integrin interaction at the plasma membrane is translated into intracellular activation of STAT3 remains unresolved. Insights gained from delineating this mechanism would not only further our understanding of how non-canonical activators of STAT3 can activate this pathway, but also define how intracellular hubs regulate keratinocyte stem cell fate.

## Result

### F-Spondin domain of Mindin interacts with CD11b and activates STAT3 in keratinocytes

The nature and outcome of signal transduction triggered by protein–protein interactions are closely governed by the specific domains involved. Domains involved in the interaction not only determine the binding specificity but also influence their dynamics in space and time. Hence, to probe the mechanism by which Mindin activates STAT3 in an α-M integrin (CD11b) dependent manner, we first focused on characterising the specific domains involved in the ligand-receptor interaction. Mindin contains F-Spondin domain at the N-terminal and Thrombospondin Type-1 Repeat (TSR) at the C-terminal region (*14*). The association of integrins with their ligands often requires the presence of divalent metal ions, such as calcium (Ca2+) or magnesium (Mg2+). This metal ion-dependent interaction occurs at specific binding sites within the integrin and ligand molecules called MIDAS. Structural analysis of Mindin revealed that F-Spondin domain of Mindin contains amino acid residues that can coordinate to bind metal ions. Furthermore, it has shown that both full-length Mindin and the F-Spondin domain bind equally to CD11b (*14*). These led us to hypothesise if the F-Spondin domain of Mindin mediates CD11b driven STAT3 activation in keratinocytes. To address this, we first evaluated if F-Spondin domain is sufficient to activate STAT3 in keratinocytes. We observed that F-Spondin treatment resulted in elevated levels of pSTAT3, the active form of STAT3, compared to control treatment and the increase is comparable to that elicited by the treatment with full length Mindin (Figure 1A). Further, co-immunoprecipitation of CD11b from keratinocytes treated with purified recombinant F-Spondin showed that CD11b binds to F-Spondin (Supplementary Figure1A). We also evaluated if the interaction between F-Spondin and CD11b on keratinocytes is MIDAS dependent. For this, we generated a recombinant mutant protein, wherein, a critical MIDAS forming residue, glutamate, at position 115 in the F-Spondin domain was changed to alanine. Co-Immunoprecipitation data revealed that mutant protein, F-Spondin (E115A), failed to interact to CD11b in keratinocytes (Supplementary Figure1A). Based on the interaction data, we investigated whether F-Spondin mediated STAT3 activation is dependent on its interaction with CD11b. WT keratinocytes treated with F-Spondin (E115A) mutant, which resulted in loss of interaction with CD11b, failed to active STAT3 (Figure 1B). Additionally, blocking ligand interaction with an inhibitory antibody against CD11b (*15*), also resulted in the failure to activate STAT3 even in the presence of F-spondin (Figure 1C). These findings establish the F-Spondin domain of Mindin to be the minimal domain required for interaction with the CD11b receptor and to initiate STAT3 signalling in keratinocytes.

**Fig. 1.**
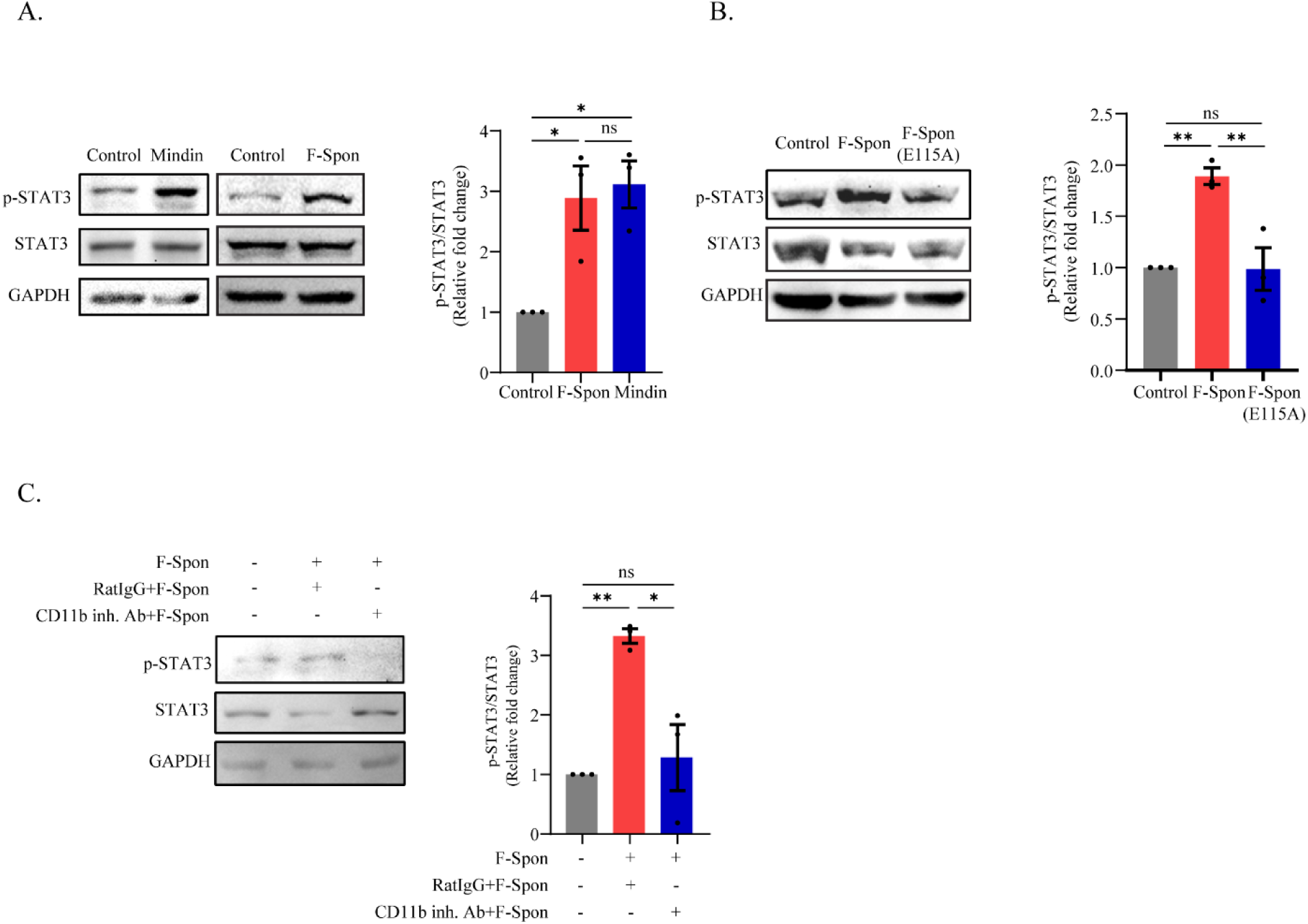
The F-Spondin domain of Mindin binds CD11b and activates STAT3 in primary mouse keratinocytes. **(A)** Left: Immunoblot; Right: Quantification of phosphorylated STAT3 (p-STAT3) relative to total STAT3 in keratinocytes treated with recombinant Mindin or F-Spondin (F-Spon) (n=3). **(B)** Left: Immunoblot; Right: densitometry of p-STAT3/STAT3 in cells treated with wild-type F-Spon or mutant F-Spon(E115A) (n=3). **(C)** Left: Immunoblot; Right: densitometry of p-STAT3/STAT3 in cells treated with F-Spon ± CD11b-blocking antibody (n=3). Data are mean ± SEM; Student’s t-test; *p < 0.05, **p < 0.01, ns = not significant.

### F-Spondin domain is sufficient to maintain STAT3 driven stemness in keratinocytes

Previously we found that the interaction between Mindin and CD11b maintains the stemness of keratinocytes in a STAT3-dependent fashion (*13*). Thus, we evaluated whether the F-Spondin domain of Mindin alone is sufficient to mediate this effect. Transcriptome analysis of F-Spondin treated keratinocytes revealed that almost 80% (237 out of 331, Figure 2A) of differentially expressed genes (adjusted *P* < 0.05, log_2_ fold change ≥1, or log_2_ fold change ≤ −1) overlap with that of Mindin treated keratinocytes (Figure 2A, Supplementary Figure 2A). Gene Ontology analysis of the overlapping gene set revealed significant enrichment in biological processes related to cell proliferation and differentiation (Figure 2B). Within these pathways, we identified a marked upregulation of epidermal stemness-associated drivers accompanied by a concomitant downregulation of key differentiation regulators (Supplementary Figure 2A). These transcriptomic trends were further validated by qPCR, which confirmed the upregulation of proliferation, survival, and keratinocyte-specific stemness genes in both Mindin- and F-Spondin–treated keratinocytes (Figure 2C). Consistent with this, we observed that F-Spondin treated keratinocytes exhibited higher nuclear to cytoplasmic ratio (Figure 2D and 2E), an increased colony-forming capacity (Figure 2F), and upregulation of CD44 expression, a keratinocyte stem cell specific surface marker (*16*) (Figure 2G). In order to understand the impact of these characteristics in maintaining the stemness of the epidermal keratinocytes, we investigated the effect of F-spondin in the calcium induced differentiation of these cells. Keratinocytes cultured under control conditions exhibited a robust and time-dependent downregulation of the epidermal stem cell marker Keratin-5, with a marked reduction detectable as early as 12 hours after calcium induction. In contrast, F-Spondin–treated keratinocytes maintained relatively stable Keratin-5 expression, showing a decrease only at 24 hours post the induction of differentiation (Figure 2H). On the other hand, examination of an early differentiation marker (Keratin 1) revealed that control keratinocytes entered the terminal differentiation program beginning at 12 hours post calcium switch, whereas F-Spondin–treated keratinocytes showed no significant induction of differentiation until 24 hours (Figure 2I). These results indicate that F-Spondin sustains stem cell marker expression while delaying the timely activation of differentiation markers.

**Fig. 2.**
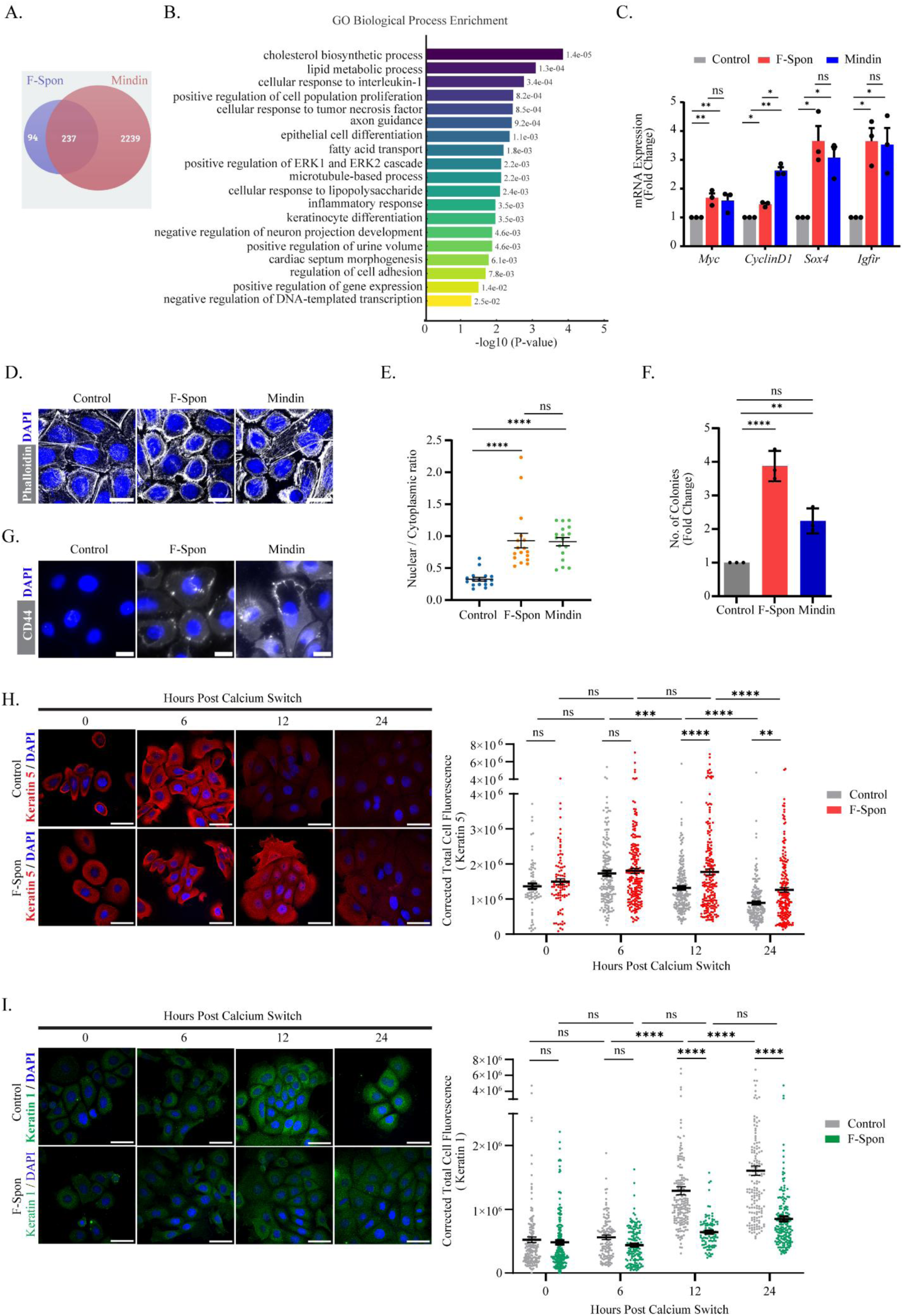
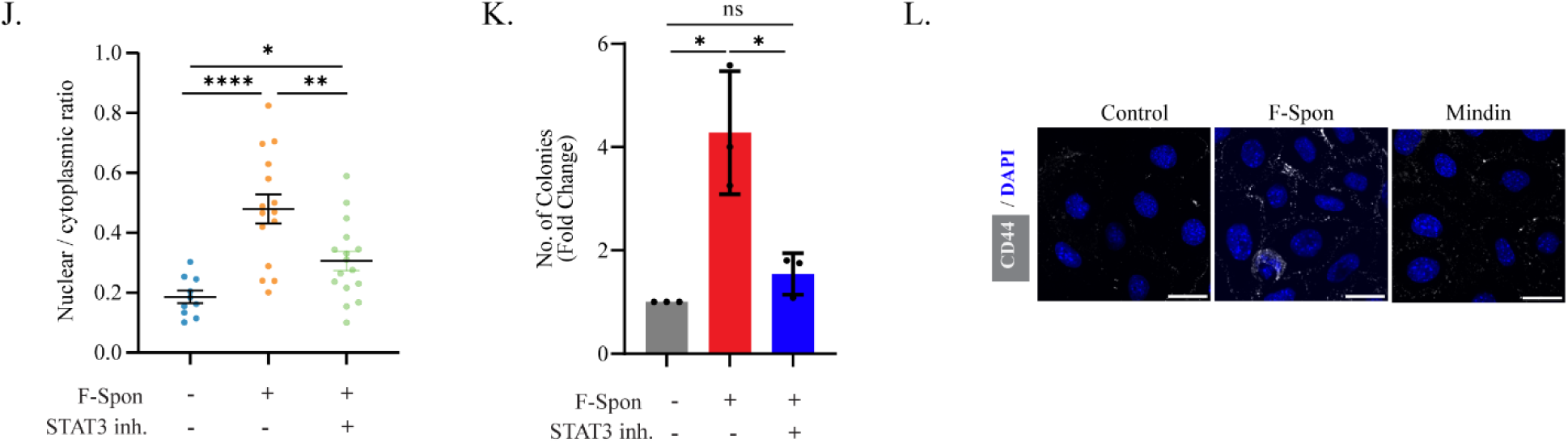
F-Spondin domain is sufficient to maintain keratinocyte stemness. **(A)** Venn diagram of overlapping differentially expressed genes in Mindin- and F-Spondin (F-Spon)–treated mouse keratinocytes (adj. P < 0.05, log2FC ≥ |1|). **(B)** Enriched GO biological processes of common genes (Fold enrichment = –log10padj, padj < 0.05). **(C)** qPCR of proliferation (*Cyclin D1*), survival (*Myc*), and stemness genes (*Sox4, Igfir*) represented as fold change. Gene expression was normalized to *TBP* (which encodes Tata Binding Protein) (n=3). **(D)** Immunofluorescence of phalloidin (actin/cell boundary) and DAPI (nucleus) in control, F-Spon, and Mindin treated keratinocytes. Scale bar: 20 µm. **(E)** Nuclear/cytoplasmic ratio from phalloidin/Hoechst images (cells from n=3 experiments). **(F)** Colony numbers in control vs F-Spon and Mindin (fold change, n=3). **(G)** Immunofluorescence of stem cell marker CD44 (gray) and DAPI in treated keratinocytes. Scale: 20 µm. **(H)** Left: Keratin-5 (basal stem/progenitor marker, red) and DAPI (blue). Scale bar: 50 µm. Right: corrected total fluorescence quantification (∼200 cells, from n=3 biological replicates). **(I)** Left: Keratin-1 (differentiation marker, green) and DAPI. Scale bar: 50 µm. Right: corrected total fluorescence (∼200 cells, n=3). **(J)** Nuclear: cytoplasmic ratio from phalloidin/DAPI images (cells from n=3 experiments). **(K)** Colony numbers in control, F-Spon, and F-Spon + STAT3 inhibitor treated keratinocytes (n=3). **(L)** CD44 (gray) and DAPI in control, F-Spon, and F-Spon + STAT3 inhibitor. Scale bar: 25 µm. Quantification data are mean ± SEM. P values: Student’s t-test (C, E, F, J, K), two-way ANOVA (H, I). *p < 0.05, **p < 0.01, ***p < 0.001, ****p < 0.0001, ns = not significant.

We further verified whether the F-Spondin domain-mediated stemness is dependent on STAT3 activation. We observed that the increase in nuclear to cytoplasmic ratio (Figure 2J), colony forming ability (Figure 2K), and CD44 expression (Figure 2L) stimulated with F-Spondin treatment was significantly blocked upon STAT3 inhibition. Altogether, these data imply that the F-Spondin domain of Mindin is sufficient to maintain STAT3-driven stemness properties of keratinocytes.

### F-Spondin mediated endocytosis of CD11b is necessary for STAT3 activation and stemness

We next investigated the mechanism by which the engagement of F-Spondin with CD11b at the cell surface results in the intracellular activation of a transcription factor. Signalling can occur at both the cell surface (such as EGF signalling) or upon internalization of the ligand-receptor complex (such as TGFβ). Flow cytometry analysis revealed that the percentage of cells expressing surface CD11b was reduced upon F-Spondin treatment (Figure 3A). However, the total CD11b protein levels are the same under both conditions (Figure S3A). This suggests that CD11b undergoes internalisation upon F-Spondin treatment. Receptor internalization in cells occurs via endocytic pathways mediated by endosomes (*17*) (*4*). Therefore, we investigated whether CD11b is redistributed to the endosome upon treatment with F-Spondin. We observed that F-Spondin treatment induced a marked colocalization of CD11b with early endosomes, marked by Early Endosome Antigen 1 (EEA-1) (Figure 3B). In the representative merged images, this colocalization was evident as prominent yellow puncta, arising from the overlap of CD11b (green) and EEA-1 (red) signals. Quantitative analysis using Pearson’s correlation coefficient (PCC), which measures the pixel-by-pixel linear correlation of fluorescence intensities between two channels, demonstrated a significant increase in the overlap between CD11b and EEA-1 upon F-Spondin stimulation compared to control conditions. Importantly, blockade of CD11b with a neutralizing antibody markedly diminished this colocalization to levels found in untreated keratinocytes (Figure 3B). Further, in ligand-receptor-mediated endocytosis, it is generally assumed that the ligand is internalized together with its receptor. We examined whether F-Spondin is endocytosed along with CD11b by triple colocalization analysis. Colocalised pixel heatmap from immunofluorescence images revealed a strong colocalization between F-Spondin-EEA1-CD11b (Figure 3C). This is further supported by the observation that the triple-colocalized pixel maps to the centre of Maxwell’s triangle, a chromaticity diagram used to represent the relative contributions of three fluorescent signals (*18*), indicating balanced signal intensities from all three channels and thereby suggesting true colocalization (Figure 3C). These data indicate that F-Spondin leads to endocytosis of the ligand-receptor complex.

**Fig. 3.**
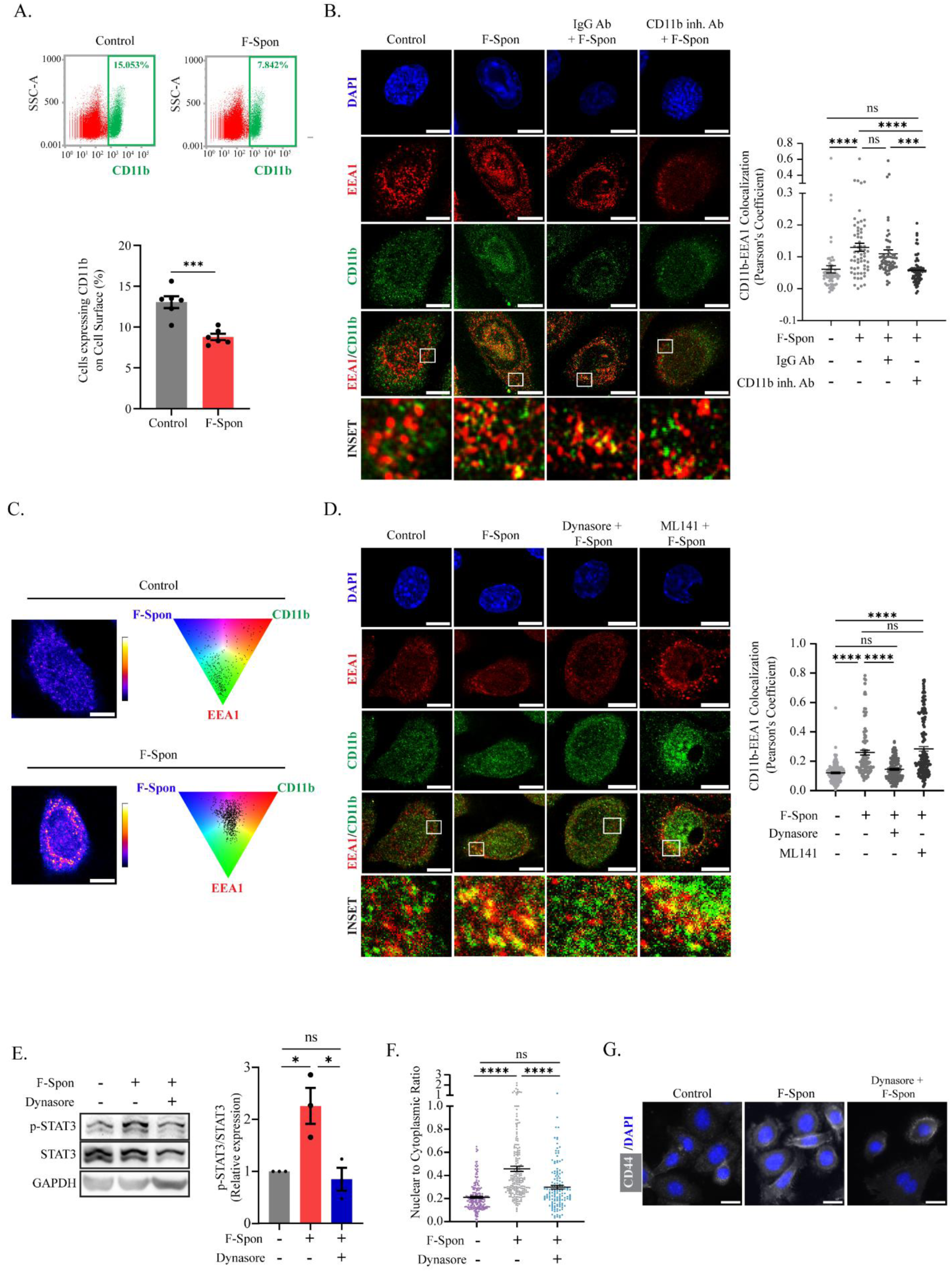
F-Spondin–mediated endocytosis of CD11b is required for STAT3 activation and stemness in keratinocytes. **(A)** Top panels: Flow cytometry of surface CD11b levels in control vs F-Spondin (F-Spon) treated keratinocytes. Percentage of cells positive(green) or negative(red) for cell surface CD11b. Quantification in the bottom panel (n = 6). **(B)** Left: Colocalization of CD11b (green) and EEA1 (red) in control, F-Spon, IgG + F-Spon, or CD11b-blocking Ab + F-Spon; DAPI stains nuclei. Scale bar: 10 µm. Right: Pearson’s correlation quantification (n = 50–80 cells from three independent experiments). **(C)** Left: Heatmap of triple colocalization (CD11b, EEA1, F-Spon) generated in ImageJ (Image Calculator); warmer colours = stronger overlap. Scale bar: 10 µm. Right: Maxwell’s triangle plots showing fluorophore contribution and overlap distribution. **(D)** Left: Colocalization of CD11b (green) and EEA1 (red) in control, F-Spon, Dynasore + F-Spon, or ML141 + F-Spon. Scale Bar: 10 µm. Right: quantification as Pearson’s correlation (n = 100–150 cells from three independent experiments). **(E)** Immunoblot of p-STAT3, STAT3, and GAPDH in control, F-Spon, and Dynasore + F-Spon with quantification (n = 3). **(F)** Nuclear-to-cytoplasmic ratio in control, F-Spon, and Dynasore + F-Spon (n = 80–100 cells). (G) CD44 (gray)/Hoechst staining under control, F-Spon, and Dynasore + F-Spon. Scale bar: 25 µm. Data = mean ± SEM. Statistics: Student’s t-test (A, E, F) or Kruskal–Wallis test (B, D). *p < 0.05, **p < 0.01, ***p < 0.001, ****p < 0.0001, ns = not significant.

Furthermore, we investigated the specific endocytic pathway utilized by CD11b during its internalization. The two major endocytic pathways are classified as either clathrin-dependent or clathrin-independent (*19*). Among the major clathrin-independent pathways is the CLIC/GEEC pathway. Pathway-specific Inhibitors were utilized in cells treated with F-spondin, and endocytosis of CD11b was determined by measuring the extent of colocalization of CD11b with EEA1. Inhibition of the clathrin-dependent pathway was achieved with the pharmacological inhibitor dynasore, that inactivates dynamin. Whereas cdc42 Rho GTPase activity which is crucial to the CLIC/GEEC pathway, was inhibited with the pharmacological inhibitor ML141. Colocalisation analysis revealed that dynasore treatment in the presence of F-spondin, inhibited the endocytosis of CD11b, while ML141 failed to block the endocytosis (Figure 3D). These data suggest that F-Spondin induces F-Spondin-CD11b internalisation via the clathrin mediated endocytic pathway.

We further tested whether the F-Spondin-mediated activation of STAT3 is dependent on endocytosis of the CD11b receptor. Cells treated with F-spondin in the presence of dynasore failed to activate the STAT3 pathway, as indicated by the p-STAT3 levels that were similar to buffer-treated cells (Figure 3E). On the other hand, as expected, impairment of the CLIC/GEEC pathway did not affect the ability of F-spondin to activate STAT3 (Figure S3B). We also check if the enhanced stemness properties in keratinocytes by F-Spondin are altered by blocking endocytosis. The increase in nuclear to cytoplasmic ratio mediated by F-Spondin was prevented upon blocking endocytosis (Figure 3F). Similarly, we did not observe CD44 expression in keratinocytes treated with dynasore (Figure 3G). Altogether, these data reveal that the F-Spondin-mediated CD11b endocytosis is necessary for STAT3 activation and associated stemness in keratinocytes.

### F-Spondin mediated endocytosis of CD11b is dependent upon Src kinase activation

We previously reported that the interaction between Mindin and CD11b activates Src kinase which mediates the downstream activation of STAT3 in keratinocytes (*13*). However, the precise role of Src kinase in this process remained unclear. Ligand-receptor interaction mediated Src kinase activation has been previously implicated in the regulation of clathrin-mediated endocytosis (*20–22*). We therefore hypothesized that Src kinase regulates STAT3 activation by driving CD11b endocytosis. To test this, we first evaluated if F-Spondin domain of Mindin is sufficient to activate Src kinase. We observed increased levels of p-Src upon F-Spondin treatment in keratinocytes (Figure 4A). Furthermore, inhibition of Src kinase activity in F-Spondin–treated keratinocytes abolished STAT3 activation (Figure 4B). We then investigate if the kinase activity regulates CD11b endocytosis. Flow cytometry analysis revealed that inhibition of Src activity led to failure in the internalisation of CD11b upon F-Spondin treatment (Figure 4C). Similarly, colocalization of CD11b in early endosomes was also blocked upon Src kinase activity inhibition (Figure 4D). In summary, these results suggest that Src kinase couples F-Spondin–CD11b interaction to the endocytosis of CD11b and subsequent STAT3 activation. Finally, we evaluated if Src activity contributes to the maintenance of stemness properties induced by F-Spondin in keratinocytes. As expected, F-Spondin mediated higher nuclear to cytoplasmic ratio (Figure 4E) and CD44 expression (Figure 4F) but this effect was inhibited upon blocking Src activity.

**Fig. 4.**
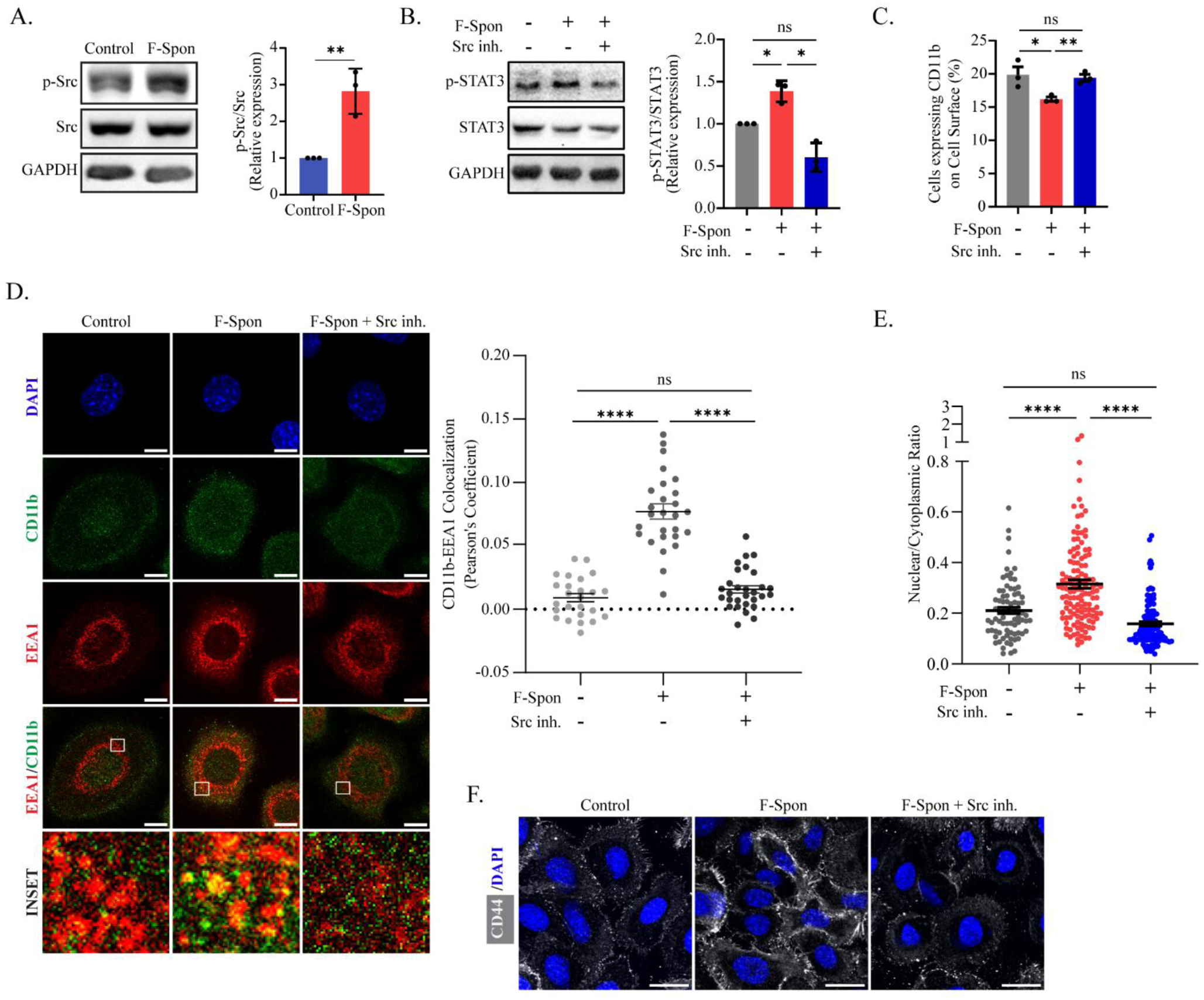
Endocytosis of CD11b is dependent upon Src kinase activity. **(A)** Left: Immunoblot of p-Src, Src and GAPDH. Right: Quantification of p-Src/Src level, n=3. **(B)** Left: Immunoblot of p-STAT3, STAT3 and GAPDH in control, F-Spon and Src inhibitor(inh.) + F-Spon treated keratinocytes. Right: Quantification of p-STAT3/STAT3, n=3. **(C)** Left: Flow cytometry data showing percentage of cells with surface CD11b levels in control, F-Spon and Src inh. + F-Spon treated keratinocytes. Right: Quantification, n=3. **(D)** Colocalization of CD11b (green) and EEA1 (red) in control, F-Spon and Src inh. + F-Spon treated keratinocytes. Scale bar: 10 µm Right: Pearson’s correlation quantification (n=20-30 cells from three independent experiments). **(E)** Nuclear to cytoplasmic ratio in control, F-Spon and Src inh. + F-Spon treated keratinocytes (n=80-100 cells from three independent experiments). **(F)** CD44 (gray)/Hoechst staining under control, F-Spon and Src inh. + F-Spon. Scale bar: 25 µm. Data represent the mean ± SEM. p values calculated using student’s t-test (A-C, E) and Kruskal-Wallis test(D), *p < 0.05, **p < 0.01, ***p <0.001 and ****p <0.0001 and ns = not significant.

### The acidic environment of endosomes induces conformational changes in CD11b–CD18 that promotes STAT3 activation

Our results demonstrates that the F-Spondin–CD11b interaction promotes Src kinase–dependent endocytosis of the receptor, which subsequently drives STAT3 activation in keratinocytes. Although endosomes have been recognized as platforms that activates signalling pathways, the mechanistic basis for this remains undefined. To address what mediates integrin dependent STAT3 activation at the endosomes, we first evaluated to what extent does ligand-receptor interaction affect integrin activation making it competent for downstream signalling. Integrin activation and signal propagation is tightly regulated by conformational rearrangements from a bent, low-affinity state, to an extended closed intermediate state and finally to an extended open high-affinity conformation (*23*) (*24*). Ligand binding is known to induce conformational changes that initiate outside-in signalling in integrins. Accordingly, we examined the conformational changes induced in the CD11b–CD18 heterodimer upon interaction with F-Spondin by modelling and simulation. Given that the Protein Data Bank (PDB) provides structural information solely for the extracellular domains of the heterodimer, the full-length proteins were modelled using AlphaFold (Supplementary Figure 4A). The AlphaFold-predicted 3D model of the extracellular domain aligned closely with the available extracellular PDB structure (Supplementary Figure 4B). This supports the reliability of the AlphaFold generated models for transmembrane and cytoplasmic domains lacking experimental structures. As the CD11b-CD18 heterodimer is a transmembrane receptor, we incorporated a membrane system with an appropriate solvent at pH 7.4 (Supplementary Figure 4C). To investigate the structural impact of F-Spondin binding, we modelled a complex of the F-Spondin domain (derived from PDB) bound to the CD11b–CD18 heterodimer (Figure 5A). Molecular dynamics simulation was performed for 300 ns, and trajectory clustering using Desmond’s tool was used to extract the most representative conformational state (Figure 5B). The predominant state revealed a partial opening of the extracellular headpiece, indicative of initial rearrangement toward the active integrin ectodomain conformation. Notably, despite these extracellular changes, the α- and β-subunit cytoplasmic tails remained clasped near the transmembrane region (Figure 5C). α/β cytoplasmic tail separation is critical to accommodate interaction with cytoplasmic signalling partners and for robust integrin signal transduction (*25*). Since F-Spondin alone is insufficient to induce cytoplasmic tail separation, we hypothesized that the endosomal microenvironment would impact the availability of cytoplasmic binding motifs and thereby regulate its downstream signalling effect. A defining feature of the endosome is its acidic environment and it is known that low pH induces structural rearrangements in endocytosed proteins that affect ligand release and receptor recycling (*26*). We investigated the effect of acidic pH conditions—pH 6.0 (early endosomes) and pH 5.0 (late endosomes/lysosomes) on the conformation of the CD11b–CD18 heterodimer, with respect to the inactive conformation of the heterodimer at the cytosolic pH 7.4. To examine how pH affects the conformation of CD11b-CD18 heterodimer, we analysed the MD simulation data with Desmond’s clustering tool. This trajectory analysis yielded the representative frame that captured the dynamic changes during the 330 ns simulation trajectory under the three pH conditions: pH 7.4, pH 6.0, and pH 5.0 (Figure 5D). Cluster analysis revealed pH-dependent conformational shifts in CD11b–CD18. At pH 7.4, the integrin adopts a bent conformation with a closed headpiece (Figure 5D) and clasped cytoplasmic tails (Figure 5D and 5E), consistent with the canonical inactive state (REF). At pH 6.0, the headpiece becomes more open (Figure 5D), the transmembrane helices tilt apart (Figure 5D), and the cytoplasmic tails separate (Figure 5D and 5E), indicative of a transition toward the active conformation. At pH 5.0, however, the extracellular region appears less open than at pH 6.0 (Figure 5D) and the cytoplasmic tails are engaged (Figure 5D and 5E), suggesting a conformation closer to inactive-like state (Figure 5C). Together, these findings suggest that F-Spondin primes CD11b–CD18 for activation, while the acidic endosomal environment drives the receptor into a state necessary for downstream signalling.

**Fig. 5.**
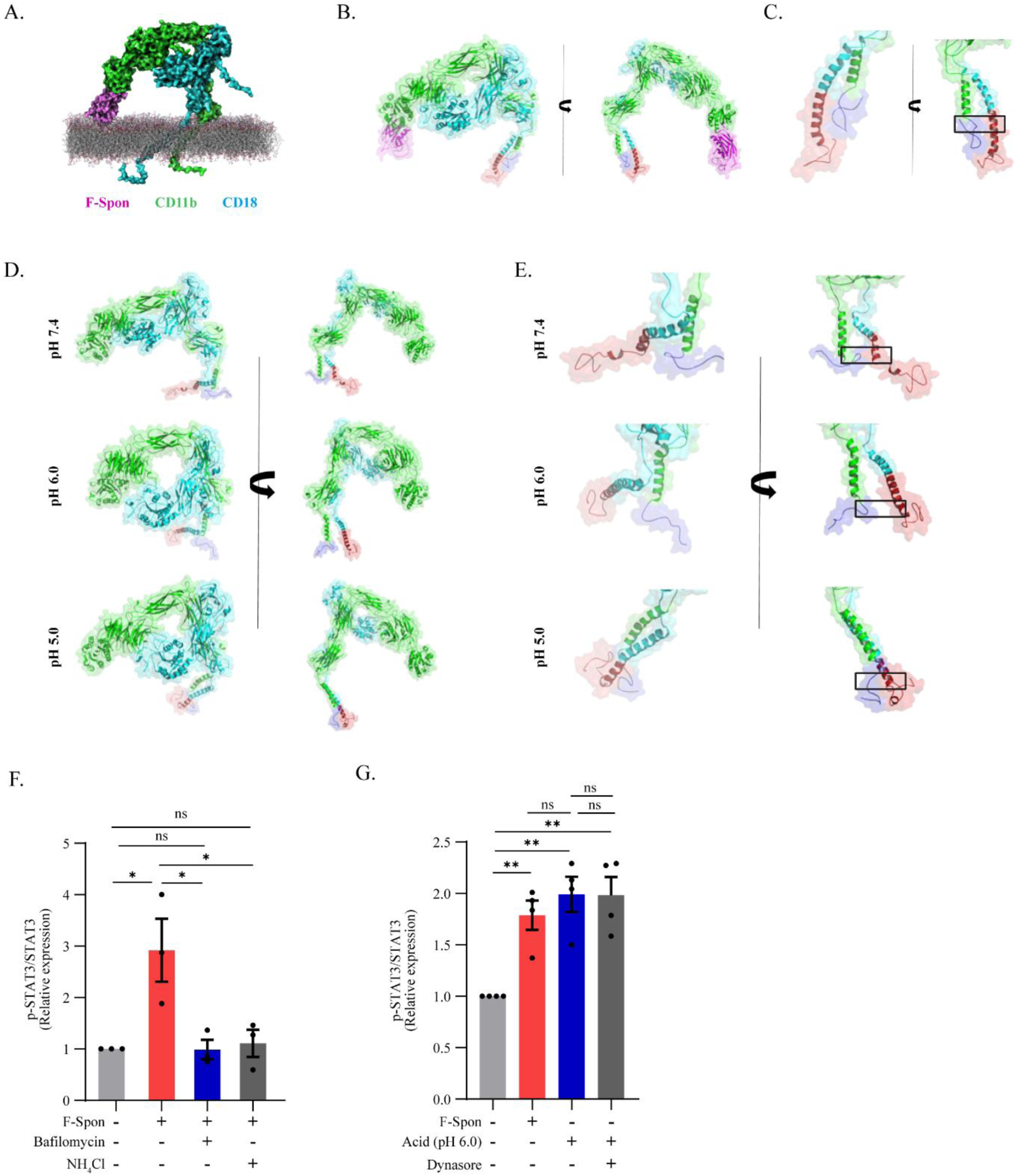
: Structural modelling of the CD11b–CD18 integrin heterodimer and regulation of STAT3 activation by endosomal acidity. (**A)** Model of the initial F-Spondin (pink)–CD11b (green)–CD18 (blue) complex generated by protein–protein docking, in which F-Spondin was aligned to high-confidence poses at the CD11b αI-domain and the heterodimer was subsequently embedded into a membrane bilayer for simulation (**B**) Surface-over-ribbon rendering of representative MD-derived clusters of the CD11b–CD18–F-Spondin complex, illustrating key conformational differences and dynamic rearrangements. (**C**) Structure of the cytoplasmic tail region in the representative MD-derived CD11b–CD18 complex. The CD11b cytoplasmic tail (Chain A, residues 1129–1152) is shown in purple, and the CD18 cytoplasmic tail (Chain B, residues 724–769) is shown in red. The black box identifies the tail–tail interaction site that governs integrin activation and is required for propagating intracellular signalling events. (**D**) Surface-over-ribbon rendering of representative MD-derived clusters of the CD11b–CD18 heterodimer simulated under different pH conditions (7.4, 6.0, and 5.0), highlighting pH-dependent conformational transitions. **(E)** Cytoplasmic tail region of representative MD-derived CD11b–CD18 complexes under varying pH conditions. (**F**) Quantification of phosphorylated STAT3 (p-STAT3) relative to total STAT3 in keratinocytes treated with F-Spondin, F-Spondin ± Bafilomycin, or F-Spondin ± NH₄Cl (n = 3). (**G**) Quantification of p-STAT3 relative to total STAT3 in STAT3-overexpressing keratinocytes treated with F-Spondin, acidic media (pH 6.0), or acidic media ± Dynasore (n = 3). Data represent mean ± SEM; Student’s t-test; *p < 0.05, **p < 0.01, ns = not significant.

Based on these models, we tested the effect of acidic pH on STAT3 activation in our in-vitro primary keratinocyte culture system. Treatment of keratinocytes with Bafilomycin A1, a well-known inhibitor of the vacuolar H⁺-ATPase (V-ATPase) present in endosomal vesicles, resulted in the failure of F-Spondin induced STAT3 activation (Figure 5F and Supplementary Figure 4D). Similarly, treatment with a weak base, NH₄Cl, which diffuses into acidic compartments and raises the pH by buffering the acidity, also blocked STAT3 activation despite the presence of F-Spondin (Figure 5F and Supplementary Figure 4D). This supports the model that low pH, maintained in the endosomal vesicle, is necessary for STAT3 activation. We tested the sufficiency of acidic pH in inducing STAT3 activation, by exposing the whole cell to the acidic pH condition of the early endosome in the absence of F-Spondin. Similar to F-Spondin treatment, exposure to acidic media led to STAT3 activation (Figure 5G and Supplementary Figure 4E). Blocking endocytosis with dynasore in the presence of acidic media did not inhibit STAT3 activation. These data demonstrates that the acidic environment of early endosomes induces conformation changes in the integrin cytoplasmic tail that facilitates the activation of STAT3.

## Discussion

Our study reveals a previously unrecognized spatial mechanism by which the extracellular matrix protein Mindin orchestrates CD11b–CD18 integrin trafficking, Src kinase activation, and STAT3 signalling to maintain keratinocyte stemness (Figure 6). We have identified the F-Spondin domain of Mindin as the minimal region sufficient for αM-integrin engagement and STAT3-driven stemness. Mechanistically, F-Spondin binding to CD11b promotes its internalization from the plasma membrane to early endosomes through clathrin-mediated endocytosis. The process of CD11b endocytosis is controlled by Src kinase activation. It has been previously reported that Src kinase facilitates clathrin-mediated endocytosis by phosphorylating components of the endocytic machinery, including clathrin heavy chain and adaptor proteins such as epsin and dynamin. These phosphorylation events promote clathrin coat recruitment and vesicle internalization, linking Src activity to receptor trafficking (*20–22*). Notably, we observed that endocytic trafficking of integrin is necessary for STAT3 activation within endosomes. The role of Mindin mediated integrin endocytosis aligns with increasing evidence linking endocytosis to the regulation of different stem cell systems. In embryonic and haematopoietic stem cells endosomal scaffolding protein OCIAD1 anchors STAT3 at early endosomes, maintaining pluripotency(*10*). Similarly, in intestinal stem cells, endocytic regulation of Wnt receptors controls β-catenin signalling strength and stem cell proliferation (*27*).

**Fig. 6:**
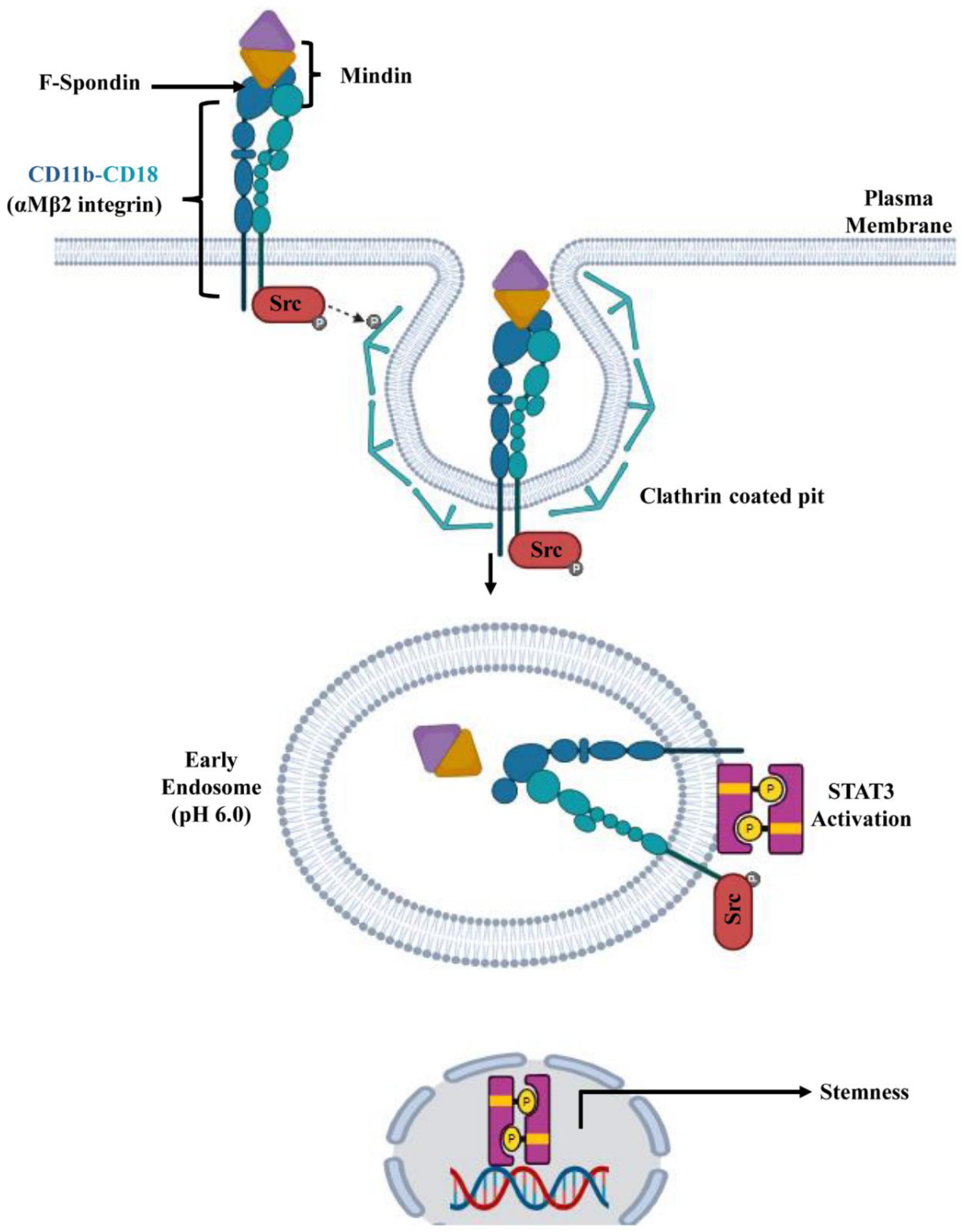
Model. Schematic showing Mindin–αMβ2 engagement triggers Src-dependent endocytosis, followed by integrin activation within the acidic endosomal compartment, which in turn drives STAT3 activation to sustain keratinocyte stemness.

A previous report suggests that the full activation of integrins at the plasma membrane can be sustained by its relocalisation to the endosomes (*28*). In the context of Mindin signalling, our data reveals that integrins undergo partial activation at the plasma membrane, and full activation (in relation to downstream signalling) is achieved upon endocytosis and trafficking to the endosome. This is demonstrated by our modelling and simulation analyses, which suggest that the F-Spondin–CD11b–CD18 interaction alone is insufficient to induce a fully active, signalling-competent integrin state (*24*, *25*, *29*, *30*). In this state, the headpiece is partially open while the cytoplasmic tails remain clasped together. Full activation, characterized by both headpiece opening and tail separation, is ultimately achieved within the acidic microenvironment of endosomes. We further validate this model experimentally by demonstrating that neutralization of endosomal pH abolished STAT3 activation. Moreover, exposure of cells to an acidic extracellular milieu was sufficient to induce STAT3 activation, when integrins were still localised at the plasma membrane. These findings demonstrate that ligand engagement primes integrin for internalization, and endosomal acidity subsequently activates integrin to trigger STAT3 activation. Previous reports have shown that extracellular acidosis in the tumour microenvironment modulates signalling proteins to regulate intracellular pathways, thereby promoting cell survival and invasive behaviour. (*31*, *32*). For instance, acidic pH has been reported to modulate the structure and function of integrins such as α5β1 and αvβ3 by inducing conformational changes toward a high-affinity active state, which, in turn, drives cancer cell migration (*33*, *34*). These observations are consistent with our data showing that exposure to pH 6.0 induces conformational changes in the CD11b–CD18 integrin that more closely resemble the active state than that induced by F-Spondin binding alone. Together, these results underscore the importance of biochemical cues, such as acidity, in fine-tuning signal transduction. However, the precise mechanism by which integrin activation in the acidic milieu of endosomes links to STAT3 activation remains an open question.

Given that the acidic tumour microenvironment promotes cell migration and invasion during tumour progression, we hypothesized that acidity may likewise regulate oncogenic signalling in cancer stem cells, the key drivers of tumour growth and recurrence. Previously, our lab developed a transgenic mouse, wherein, the transcription factor Snail is overexpressed in the keratinocyte stem cells present in the basal layer of the skin epidermis. This system has been shown to recapitulate key hallmarks of squamous cell carcinoma including epidermal hyperplasia (*35*). Snail-transgenic keratinocytes exhibit cancer stem cell like properties, and our earlier work demonstrated that Mindin regulates CD11b-dependent activation of STAT3 in these cells to sustain stemness programs and drive epidermal hyperplasia (*13*). Given that Mindin-CD11b mediated STAT3 activation depends on CD11b trafficking to acidic endosomes, we tested if this step regulates oncogenic STAT3 activation and stemness in this cancer stem cell model. We observed reduced levels of CD11b at the cell surface (Supplementary Figure 5A) and increased accumulation of CD11b within endosomes (Supplementary Figure 5B) in Snail-transgenic keratinocytes compared to wild-type cells. Blocking CD11b endocytosis in Snail transgenic keratinocytes reversed this distribution, restoring surface expression to wild-type levels and eliminating endosomal accumulation (Supplementary Figure 5A and Figure 5B). Notably, inhibition of endocytosis abolished STAT3 phosphorylation (Supplementary Figure 5C) and reduced features of stemness (Supplementary Figure 5D and Figure 5E) in Snail transgenic keratinocytes. Together, these findings establish that trafficking of CD11b to the endosomes that exposes it to an acidic microenvironment serves as a critical regulator for oncogenic STAT3 activation and stemness maintenance in a model of cutaneous carcinoma.

Consistent with our findings, it is increasingly evident that endocytosis is a key regulator of oncogenic signalling by controlling receptor availability, signal duration, and subcellular localization across multiple cancers (*12*). Endocytosis inhibitors such as Dynasore, MiTMAB, and Dynole-34-2 have demonstrated potent anti-proliferative and pro-apoptotic effects in cervical cancer and other tumour cell lines, suggesting their potential in anti-cancer treatments (*36*). Given our findings that endocytosis-driven signalling maintains stemness in an epidermal cancer stem cell model, targeting this pathway with endocytic inhibitors may represent a promising therapeutic strategy.

## Materials and Methods

### Cell Culture

Primary epidermal keratinocytes were isolated from respective wild type and Snail transgenic B6J-2019 mouse strains and cultured as described previously(*37*). The keratinocytes were grown in low calcium (50uM) E-media. For differentiation assay, E-media containing 1.2mM calcium was used and kept for indicated time points.

### Antibodies and Reagents

The following antibodies were used in the study: p-STAT3 (9145S-CST), STAT3 (4904-CST), GAPDH (ab8245-abcam), p-Src (2101-CST), Src (2123-CST), CD11b (14-0112-82-eBiosciences for immunofluorescence and immunoprecipitation, 50-161-42-Invitrogen™ for flow cytometry and PA5-90724-Invitrogen for western blot), RatIgG isotype control (12-432180-Invitrogen), EEA1 (BD610456 BD biosciences), F-Spondin (HPA066095-Sigma), CD44 (550538-BD). Keratin-1 and Keratin-5 (Jamora lab generated).

The following reagents were used: Hoechst (H3570-Life Technologies), Dynasore (D7693-Sigma-Aldrich), Src inhibitor (4660-Tocris), STAT3 inhibitor (573132-Millipore), Bafilomycin A1 (19-148-Sigma-Aldrich), ML141 (HY-12755-MedChemExpress)

### Protein purification

The cDNA encoding the F-Spondin domain of Mindin was cloned into a pET-28a expression vector with an N-terminal 6×His tag and transformed into *E. coli* BL21 (DE3) competent cells. A single colony was inoculated into LB medium supplemented with kanamycin and grown overnight at 37 °C. The overnight culture was diluted 1:100 into fresh LB medium and grown at 37 °C until OD₆₀₀ reached 0.6–0.8, followed by induction with 250 mM IPTG at 18 °C for 10-12 hrs with constant shaking at 180rpm.

Cells were harvested by centrifugation (5000 × g, 15 min, 4 °C), resuspended in lysis buffer (20 mM Tris-HCl pH 8.0, 300 mM NaCl, 10 mM imidazole, 1 mM PMSF), and lysed by sonication on ice. The lysate was clarified by centrifugation at 15,000 × g for 45 min at 4 °C, and the supernatant was loaded onto Ni-NTA column (Biorad) pre-equilibrated with lysis buffer. The resin was washed extensively with wash buffer (20 mM Tris-HCl pH 8.0, 300 mM NaCl, 30–40 mM imidazole) to remove non-specifically bound proteins, and bound proteins were eluted using elution buffer (20 mM Tris-HCl pH 8.0, 300 mM NaCl, 100-250 mM imidazole).

The eluted protein was concentrated using Amicon Ultra centrifugal filters (Millipore, 3 kDa cutoff) and subjected to size-exclusion chromatography (SEC) on a Superdex 200 Increase 10/300 GL column (Cytiva) pre-equilibrated with SEC buffer (10 mM Tris-HCl pH 8.0, 20 mM NaCl) using an ÄKTA pure system. Peak fractions corresponding to the target protein were pooled and concentrated. Protein purity was assessed by SDS–PAGE followed by Silver-nitrate staining, and concentrations were determined using nano-drop. The purified F-Spondin was used at a concentration of 250ng/ml in all treatments. All treatments were done in serum-free conditions unless otherwise stated.

Histidine-tagged Mindin was purified from conditioned media collected from Mindin expressing Chinese Hamster Ovary cells using Ni-NTA column. Buffers with varying strengths of imidazole were made in 20 mM Tris and 300 mM NaCl (pH = 8) for washing and elution. The purified Mindin (10 mL) was dialyzed for 3 rounds in 1 L dialysis buffer (10 mM tris, 20 mM NaCl, pH = 8), concentrated using a 10 kDa Centricon concentrator, and filtered with a 0.2 μm syringe filter. Silver staining was performed to assess the purity and confirm the purification of Mindin. The purified recombinant Mindin was used at a concentration of 80–200 ng/mL in all treatments. All treatments were done in serum-free conditions unless otherwise stated.

### Western Blotting

Keratinocyte lysates were prepared in RIPA buffer (50 mM Tris-HCl, pH 7.4; 150 mM NaCl; 1% NP-40; 0.1% SDS; 0.5% sodium deoxycholate) supplemented with protease (S8830, Sigma) and phosphatase (04 906 837 001, Roche) inhibitors and sonicated on ice at 4 °C. Equal amounts of protein were mixed with 4× Laemmli sample buffer, boiled at 95 °C for 3 min, and resolved by SDS–PAGE. Proteins were transferred onto nitrocellulose membranes using a wet transfer system. Membranes were blocked for 1 h at room temperature with either 5% BSA in Tris-buffered saline containing 0.1% Tween-20 (TBST) for phosphorylated proteins, or 5% Blotto (Santa Cruz Biotechnology, sc-2325) for total proteins. Membranes were then incubated overnight at 4 °C with the appropriate primary antibodies, followed by HRP-conjugated secondary antibodies for 1 h at room temperature. After extensive washing with TBST, signals were detected using enhanced chemiluminescence (ECL, Merck) and visualized on an iBright FL (Thermo Fisher) imaging system. Band intensities were quantified using Fiji (ImageJ), and values were normalized to the respective loading controls.

### Gene expression

Total RNA was extracted from keratinocytes using RNAIso Plus (9109-Takara) according to the manufacturer’s instructions. RNA purity and concentration were assessed using a NanoDrop spectrophotometer and RNA integrity was verified by agarose gel electrophoresis. For cDNA synthesis, 2 µg of total RNA was reverse transcribed using the PrimeScript™ RT Reagent Kit (Perfect Real Time, RR037A, Takara) following the recommended protocol.

Quantitative real-time PCR (qRT-PCR) was performed using TB Green® Premix Ex Taq™ II (Tli RNaseH Plus, RR820A, Takara). Each reaction was run in triplicate, and relative gene expression was calculated using the 2^(-ΔΔCt) method, normalized to TBP as the endogenous control.

Following is the list of primer pairs used for gene expression analysis:

*Myc:* Fwd-CCTCACTCCTAATCCGGTCAT

Rvs-GTGCTGTAGTTTTTCGTTCACTG

*CyclinD1*: Fwd-GCGTACCCTGACACCAATCTC

Rvs-CTCCTCTTCGCACTTCTGCTC

*Sox4*: Fwd-GACAGCGACAAGATTCCGTTC

Rvs-GTTGCCCGACTTCACCTTC

*IGIF1R*: Fwd-GTGGGGGCTCGTGTTTCTC

Rvs-GATCACCGTGCAGTTTTCCA

*TBP*: Fwd-AGTGCCGCCCAAGTAGCA

Rvs-TCCCCCTCTGCACGTAAATC

### Immunoprecipitation

Keratinocytes were lysed in ice-cold lysis buffer (50 mM Tris-HCl, pH 7.4; 150 mM NaCl; 1% NP-40) supplemented with protease (S8830, Sigma) and phosphatase (04 906 837 001, Roche) inhibitors. For each reaction, equal amounts of protein were incubated overnight at 4 °C with the indicated primary antibody or isotype control IgG. Antibody–protein complexes were captured by incubation with Protein G agarose beads (Thermo Fisher Scientific) for 30 mins. at room temperature with gentle rotation. Beads were washed three times with lysis buffer, and bound proteins were eluted by boiling in 2× Laemmli sample buffer for 10 min at 95 °C. Eluates were resolved by SDS–PAGE and analyzed by immunoblotting with the respective antibodies.

### Immunostaining and image acquisition

Primary mouse keratinocytes were seeded on 10 mm circular coverslips coated with Rat tail Type-I collagen (50 ug/ml, Gibco-A1048301). At the end of the experiments, cells on coverslips were washed and fixed using 4%Paraformaldehyde at 25^0^C for 10 mins. The cells were then permeabilized using 0.2% Triton-X100 at 25^0^C for 10 mins. The cells were blocked using 5% BSA blocking solution in TBST for 30-60 mins. This was followed by primary antibody staining overnight at 4^0^C. The next day, primary antibody was removed followed by three washes with TBST, 10 mins. each. Cells on coverslip were then incubated with respective secondary antibodies and Hoechst (to stain the nucleus) for 30 mins. – 1 hour followed by three washes TBST for 5-7 min. each and mounted with Mowiol 4-88 (81381-50G, Sigma-Aldrich).

The images were acquired using an Olympus FV3000 confocal microscope and processed using FIJI software. Z-stack images were obtained at a resolution of 800 × 800 pixels and were subsequently visualized and generated using FIJI software.

### Colocalisation analysis

For co-localization of CD11b and EEA1, cells were imaged using a (Olympus FV3000 confocal microscope with PLAPON 60X objective having a 1.42 numerical aperture, oil immersion, and a 1 μm pinhole setting). Images were acquired as z-stacks with an optical section thickness of ∼0.3 µm to capture the full cellular volume. Co-localization analysis was performed on single optical sections to avoid artifacts from projected planes. Background fluorescence was subtracted uniformly from all channels prior to analysis. Individual cells were selected as regions of interest (ROIs) to exclude adjacent cells or extracellular signals. The degree of co-localization was quantified using the Pearson’s correlation coefficient, measured with the *Coloc2* plug-in in FIJI (ImageJ). Identical thresholding parameters were applied across datasets to minimize bias. Pearson’s coefficients were calculated per ROI, and values from individual cells were plotted as single data points in the graphs.

Triple colocalization between CD11b, EEA1 and F-Spondin was quantified using the ColocQuant/ColocJ pipeline (*18*).

### Flow Cytometry

Keratinocytes were harvested using EDTA solution and washed twice with ice-cold PBS containing 1% FBS (FACS buffer). For surface staining, 1 × 10⁶ cells were resuspended in FACS buffer and incubated with CD11b Monoclonal Antibody (M1/70), APC-eFluor™ 780 for 30 mins on ice washed twice in FACS buffer and resuspended in 500 µLFACS buffer. Data were acquired on the Attune NxT Acoustic Focusing Cytometer (Invitrogen).

### Transcriptome analysis (RNA-seq) using RNAlysis

Raw FASTQ files were imported into RNAlysis (3.9.2) Kallisto Paired-end RNA-seq quantification tool under FASTQ tab. The count matrix file from Kallisto quantification was imported for DESeq2 normalisation within RNAlysis. Low-abundance genes were filtered. Genes were called differentially expressed (DEGs) at adjusted p-values< 0.05 and |log2 fold-change| ≥ 1, unless otherwise indicated.

Lists of up-regulated and down-regulated DEGs (separately) were submitted to DAVID Bioinformatics Resources (v2023q4; build noted at submission). Enrichment was assessed for GOTERM_BP_DIRECT. Terms with adjusted p-value<0.05 were considered significant. Statistics were taken directly from DAVID.

Overlaps among DEG sets from Mindin and F-Spondin comparisons were computed on gene symbols. Venn diagrams were generated with Venny 2.0 using the exact DEG lists and thresholds defined above.

### Nuclear to Cytoplasmic ratio

Keratinocytes were cultured on coverslips, fixed with 4% paraformaldehyde (PFA) for 10 min at room temperature, permeabilized using 0.2% Triton X-100 in PBS, and stained with Hoechst (for nuclei) and phalloidin–Alexa Fluor 568 to delineate cell boundaries. Confocal z-stacks were acquired using a (Olympus FV3000 confocal microscope) under identical acquisition parameters for all conditions.

Image processing: Images were analyzed using FIJI (ImageJ). Nuclear ROIs were generated by automatic thresholding of the DAPI channel, followed by manual curation to exclude overlapping or fragmented nuclei. Cytoplasmic area was calculated from subtracting nuclear area from total cell area generated from phalloidin/cytoplasmic marker-defined cell boundary. The nuclear and cytoplasmic areas were computed from the segmented ROIs. The ratio was calculated as: N/C ratio (area)= Nuclear area/Cytoplasmic area.

### Colony forming assay

Keratinocytes were seeded at a low density (1,000 cells per well) in 6-well culture plates and allowed to adhere for 12 hours. After 12 hours Mindin/F-Spondin treatment was initiated and supplemented every alternate day. The cells were maintained under standard growth conditions in DMEM-F12 and 1XPen-Strep solution (A001A-100ML-HiMedia) for 10–14 days without disturbance, allowing colonies to form. At the end of the incubation period, colonies were washed gently with PBS, fixed with 4% paraformaldehyde for 15 min, and stained with 0.5% crystal violet solution in 25% methanol for 30 min. Excess stain was removed by washing with distilled water, and plates were air-dried. Colonies containing more than 10 cells were counted manually using a light microscope or automatically using FIJI (ImageJ) software.

### Structure retrieval and modelling

The complete 3D structure of the CD11b and CD18 heterodimer was not available in the PDB database. Therefore, individual predicted AlphaFold models from the UniProt database (https://www.uniprot.org) (UniProt IDs P11215 and P05107) were used (*38*, *39*). To model the heterodimer, the individual model structures were aligned with the heterodimer 3D crystal structure available in the PDB database (PDB ID: 7USL) using Schrödinger’s Maestro. Modeling of F-Spondin of Mindin protein bound with Cd11b-CD18 heterodimer carried out using protein-protein docking server i.e. HADDOCK (https://rascar.science.uu.nl/haddock2.4/) (*40*, *41*). Input for HADDOCK were, F-Spondin structure from PDB database (PDB ID: 3D34, chain A) and CD11b alpha-I domain from AlphaFold model (UniProt ID: P11215). Active residues were defined as Lys16, Glu96, Glu115 for F-Spondin and MIDAS binding site residues of alpha-I domain i.e. Asp156, Gly157, Ser158, Gly159, Ser160. Docked complex then utilized for CD11b-CD18-Spondin complex using Maestro.

The constructed heterodimer model was subjected to protein preparation using Maestro’s ‘Protein Preparation Wizard’ utility at three specified pH values i.e. 5.0, 6.0, and 7.4 in case of CD11b-CD18 heterodimer, while pH 7.4 was used for CD11b-CD18-Spondin complex. This step included assigning bond orders, adding missing hydrogens, creating disulfide bonds, forming zero-order bonds to metals, and capping terminals. Further refinement was performed to optimize hydrogen bonding, with protonation states of residues at specific conditions determined using a pKa prediction by PROPKA.

The refined structures were further optimized for the hydrogen-bonding network. This involved adjusting the orientation of hydroxyl groups, performing 180-degree flips of the amide groups of asparagine and glutamine, and modifying the ring orientation and charge state of histidine residues.

### Molecular dynamics simulation

Prior to conducting molecular dynamics (MD) simulations, the protein structure must be embedded in a periodic boundary condition, incorporating a membrane system. As the CD11b and CD18 heterodimer is a transmembrane protein, a membrane system with an appropriate solvent at the desired pH was required. To construct this setup, Desmond’s “System Builder” tool was used to design the membrane system at three distinct pH values: 5.0, 6.0, and 7.4. The transmembrane regions of CD11b (residues 1105–1128) and CD18 (residues 701–723) were used to define the membrane placement. The system was built using the TIP3P water model, an orthorhombic box with dimensions of 35.0 Å × 35.0 × 35.0 Å, and counter ions were added to ensure neutrality. The OPLS-2005 force field was applied for parameterization of the simulation systems (*42*).

The prepared systems were subjected to MD simulations using Schrödinger’s Desmond (*43*). Each production MD simulation ran for 330 ns except in case of spondin bound complex, i.e. ran for 300ns, with energy recorded every 1.2 ps and trajectory snapshots saved every 100 ps. Prior to production runs, the simulation box volume was equilibrated under NPγT conditions at 300 K and 1.01325 bar. Desmond’s standard relaxation protocol was employed for system equilibration.

During simulations, a time step of 2 fs was used. Temperature regulation was carried out using the Nose-Hoover chain thermostat with a relaxation time of 1.0 ps, while pressure was controlled using the Martyna-Tobias-Klein barostat with a 2.0 ps relaxation time in isotropic mode. A cutoff radius of 9.0 Å was set for short-range coulombic interactions. Trajectory cluster analysis was conducted to identify representative snapshots that reflect structural dynamics.

## Author contributions

Conceptualization: BD, KB, CJ; Methodology: BD, JA, TS, A. Shrivastava, KB, AH, SK, CJ; Investigation: BD, JA, TS, A. Shrivastava, AH, AD, SJ, SK, A. Singh; Formal Analysis: BD, JA, AS, TS, AH, AD; Visualization: BD, JA, AS, AD; Funding Acquisition: CJ; Supervision: CJ; Writing - Original Draft Preparation: BD, CJ; Writing - Review and Editing: BD, CJ, A. Shrivastava, A. Singh.

## Acknowledgments

The authors would like to thank members of the Jamora laboratory for critical review of the manuscript. We would like to thank Akanksha Srivastava for generating the F-Spondin clone. This work was supported by core funds from inStem and Shiv Nadar Institution of Eminence and by grants from the Department of Biotechnology of the Government of India (BT/PR8738/AGR/36/770/2013 and BT/PR32539/BRB/10/1814/2019) and the Indian Council of Medical Research (IIRPIG-2024-01-01387). Animals are maintained at the BLiSC Animal Care and Resource Center (ACRC) and is partially supported by the National Mouse Research Resource grant BT/ PR5981/MED/31/181/2012;2013-2016;2018 and 102/IFD/SAN/5003/2017-2018 from the Department of Biotechnology.

## Conflict of interest

The authors have declared that no conflict of interest exists.

## Notes

### Competing Interest Statement

The authors have declared no competing interest.

### Summary of Updates

Abstract, Introduction, Result, Discussion and Materials and Methods section updated to add new findings Figures revised to add more data panels and updated the existing ones Figure 5 is added as a new inclusion Authors added Supplementary data added

