## Supplementary_Figures for "Mindin mediated αM-Integrin endocytosis activates STAT3 to maintain keratinocyte stemness"

A

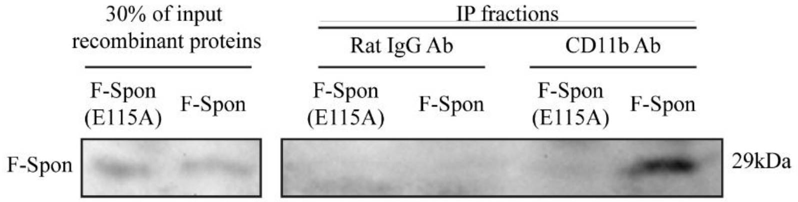


**Supplementary Fig. 1: The F-Spondin domain of Mindin binds CD11b**

**(A)** Co-immunoprecipitation of F-Spondin with CD11b from keratinocytes treated with either recombinant wild-type F-Spondin or F-Spondin mutant (FSE115A). Immunoprecipitated F-Spondin was detected by immunoblotting (Representative image among n=3 independent experiments).

A.

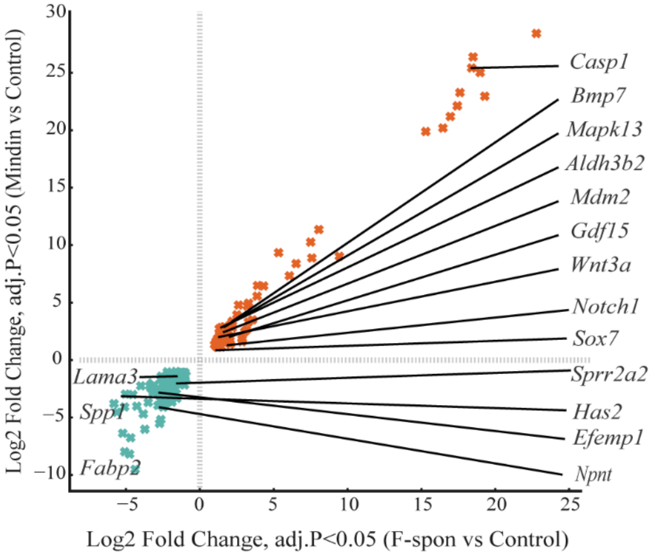

**Supplementary Fig. 2: Common regulators of proliferation and differentiation under F-Spondin and Mindin**

**(A)** Scatter plot of log2 fold changes in F-Spondin (F-Spon, x-axis) versus Mindin (y-axis) relative to control. Commonly upregulated genes (*padj* < 0.05, log2FC ≥ 1) are shown in orange and commonly downregulated genes (*padj*<0.05, log2FC≤–1) in teal. Key proliferation/stemness drivers and differentiation regulators are highlighted.

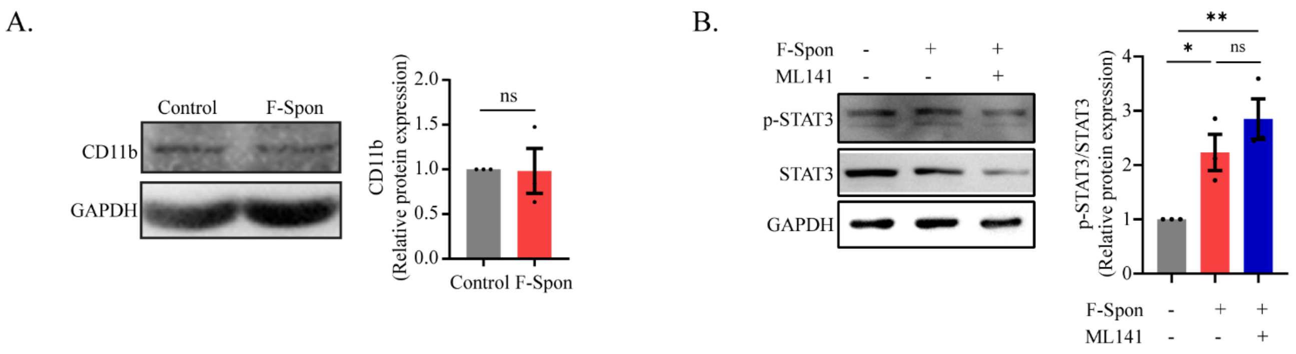


**Supplementary Fig.3: CD11b expression and STAT3 activation following F-Spondin treatment**

**(A)** Left: Immunoblot of CD11b and GAPDH (internal control) in control and F-Spondin treated keratinocytes. Right: Quantification of CD11b levels normalised to GAPDH levels from immunoblot, n=3. **(B)** Left: Representative immunoblot of p-STAT3, STAT3, and GAPDH in control, F-Spondin, and ML141+F-Spondin treated keratinocytes. Right: Quantification of p-STAT3/STAT3 levels from immunoblot, n=3. Data represent mean ± SEM; Student's t-test; *p < 0.05, **p < 0.01, ns = not significant.

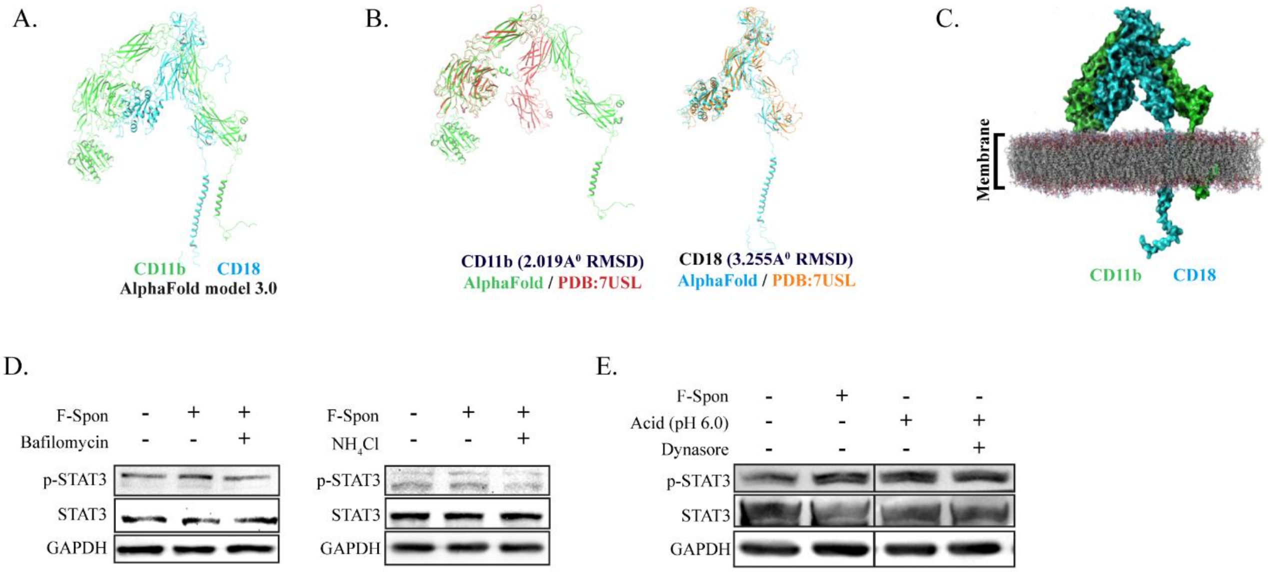


**Supplementary Fig. 4: CD11b–CD18 structure and pH control of STAT3 signalling**

**(A)** AlphaFold 3.0 prediction of the full-length CD11b (green)–CD18 (blue) heterodimer illustrating the overall architecture of the extracellular, transmembrane, and cytoplasmic regions. **(B)** Structural alignment of AlphaFold-predicted domains with the available crystal structure (PDB: 7USL). Left: CD11b (green, AlphaFold) aligned with CD11b from PDB 7USL (red), showing a backbone RMSD of 2.019 Å. Right: CD18 (blue, AlphaFold) aligned with CD18 from PDB 7USL (orange), showing a backbone RMSD of 3.255 Å. **(C)** CD11b–CD18 heterodimer embedded in a lipid bilayer, depicting the membrane positioning and spatial orientation of CD11b (green) and CD18 (blue) in a physiologically relevant environment. **(D)** Representative immunoblot of phosphorylated STAT3 (p-STAT3), total STAT3, and GAPDH in keratinocytes treated with F-Spondin, with or without inhibitors of endosomal acidification. Left: Bafilomycin treatment. Right: $NH_4Cl$ treatment. **(E)** Representative blots of p-STAT3, total STAT3, and GAPDH in STAT3-overexpressing keratinocytes treated with F-Spondin, acidic media (pH 6.0), or acidic media with the dynamin inhibitor Dynasore. The blot in this panel is shown as a split blot as demarcated by a vertical black line. One intervening lane was omitted from the final panel. All lanes were run on the same gel and imaged under identical acquisition settings.

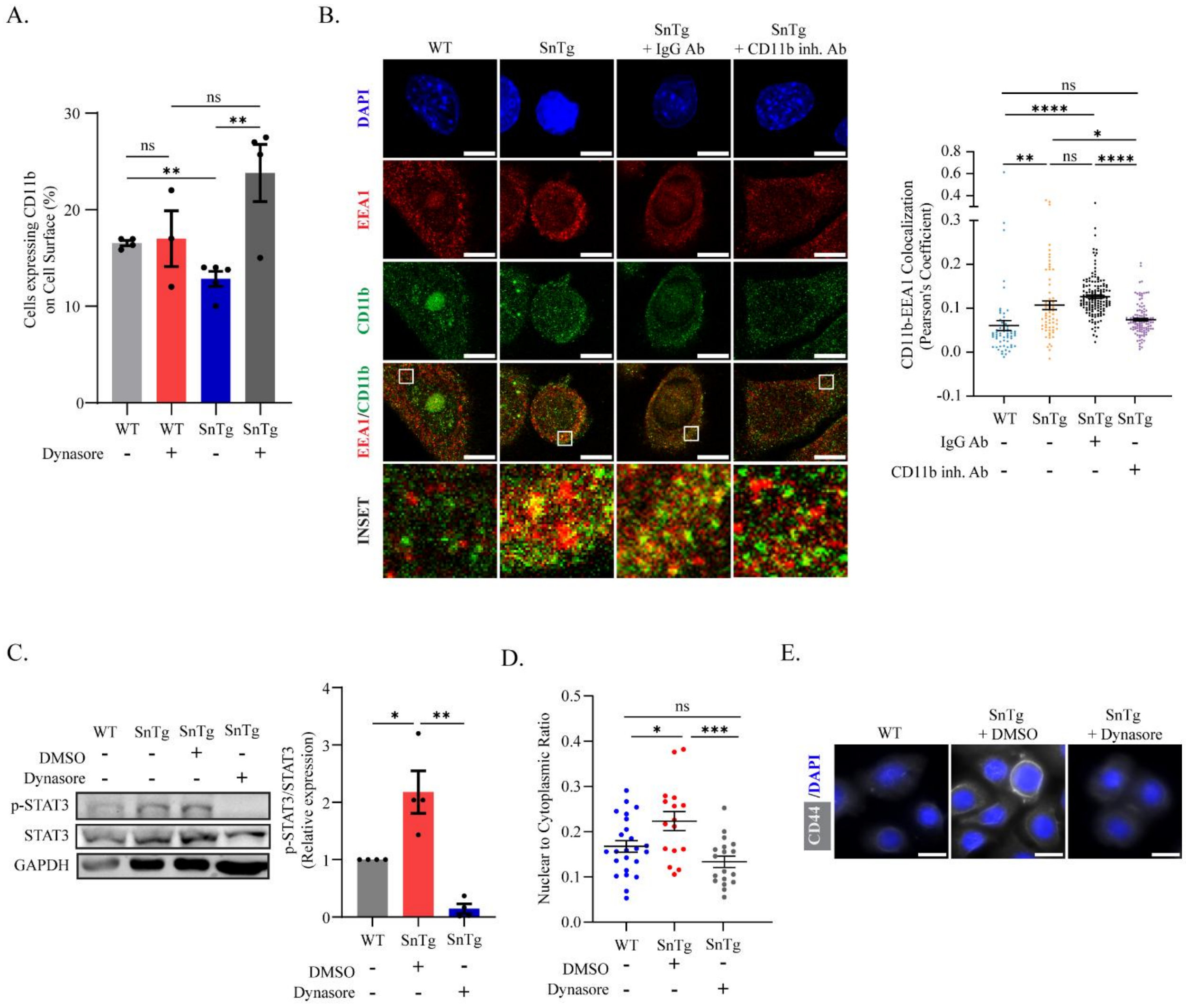


**Supplementary Fig. 5: CD11b is endocytosis in Snail transgenic keratinocytes.**

**(A)** Quantification of percentage of cells expressing CD11b surface levels in wild type (WT) primary mouse keratinocytes +/- dynasore and Snail transgenic (SnTg) mouse keratinocytes +/- dynasore, n=3-5 **(B)** Left: Representative immunofluorescence images of CD11b (in green)-EEA1(in red) colocalisation in WT and SnTg keratinocytes and SnTg treated with Rat IgG Ab and Cd11b inhibitory Ab. DAPI marks the nucleus. Images are represented from single mid-optical section. Scale Bar: 10 µm. Right: Quantification of colocalization of CD11b and EEA1 per cell in form of Pearson's correlation coefficient, n=80-100 cells from three independent experiments. **(C)** Left: Representative immunoblot of p-STAT3, STAT3 and GAPDH (internal control) in WT and SnTg keratinocytes and SnTg keratinocytes+/- dynasore. Right: Quantification of p-STAT3/STAT3 levels from immunoblot, n=3. **(D)** Quantification of Nuclear to cytoplasmic ratio in WT and SnTg keratinocytes +/- dynasore, n=30-40 cells from three independent experiments. **(E)** Immunofluorescence images of keratinocytes stained with keratinocyte stem cell marker, CD44 (in green). Hoechst marks the cell nucleus. Scale Bar: 20 µm. Data represents the mean ± SEM. p values calculated using student's t-test (A, C, D) and Kruskal-Wallis Test(B), *p < 0.05, **p < 0.01, ***p <0.001 and ****p <0.0001 and ns = not significant.
